# Nutritional Triggers of PCOS and NAFLD: A Comparative Analysis of Diets in Female C57BL/6J Mice

**DOI:** 10.1101/2025.07.21.665876

**Authors:** Sivapriya Sivagurunathan, Amit Kumar, Deepak Kumar, Meera Govindarajan, Abhiram Charan Tej Mallu, Madhulika Dixit

## Abstract

**Objective:** Obesity is a key factor in the development of Polycystic Ovary Syndrome (PCOS) and Non- alcoholic fatty liver disease (NAFLD). Its relationship with reproductive health and metabolic disorders remains an area of interest. Here we aimed to study the effects of two different obesogenic diets in female C57BL/6J mice on their metabolic and reproductive function.

**Methods:** Female C57BL/6J mice were fed high-fat high-sugar (HFHS) (45% fat and 10% sugar) diet and High fat diet (60% fat) for 21 weeks to investigate the effects on metabolic health and reproductive health. Mice were assessed for body weight, abdominal circumference, glucose tolerance (OGTT) and estrous cyclicity. At the end of the study biochemical analysis was done on isolated serum and, histological assessment of organs including liver and ovarian tissue was done to score for cystic follicle development and steatosis respectively.

**Results:** Mice on the HFHS diet did not gain weight but developed dyslipidaemia, abnormal liver enzyme levels and severe hepatic steatosis. They did not show impaired glucose tolerance and did not develop any PCOS-like ovarian morphology indicating hepatic metabolic disturbances without significant impact on reproductive health. In contrast, mice on the 60% high-fat diet (HFD) showed marginal increase in body weight and developed altered estrous cyclicity and PCOS-like ovarian morphology. They also exhibited impaired glucose tolerance and insulin resistance, confirming a metabolic disturbance associated with PCOS. These mice did not develop significant liver fibrosis, suggesting a more specific impact on the ovaries and glucose metabolism. Our findings suggest that diet composition plays a crucial role in determining the specific metabolic and reproductive outcomes in obesity-related disorders in females.

## Introduction

The global rise in obesity, particularly among women of reproductive age, presents a serious concern for both metabolic and reproductive health. Over 38.3% of reproductive aged women are classified as obese(1). Obesity is a key driver in exacerbating Insulin resistance and chronic low-grade inflammation which contribute to both ovarian dysfunction and hepatic steatosis (2–6). Obesity-related complications such as polycystic ovary syndrome (PCOS) and non-alcoholic fatty liver disease (NAFLD) are becoming increasingly prevalent. According to the World Health Organization, Polycystic ovary syndrome is one of the most common endocrine disorders in reproductively aged women, with a global prevalence of about 6-13%.(7) In India, PCOS impacts approximately 3.7 to 22.5 % of reproductively aged women depending on the population studied.(8) These women are often present with metabolic disturbances, hypertension, impaired glucose tolerance, and insulin resistance(9). In comparison to Western populations, there exists a pronounced correlation between insulin resistance and PCOS in Indian women, overshadowing the significance of hyperandrogenism(10,11).

NAFLD has been the most prevalent chronic liver disease worldwide in the past decade(12). The prevalence of NAFLD among Indian women in the general population ranges from 30% to 35%, with higher rates seen in urban areas(13). In high-risk groups such as women with PCOS, diabetes, or obesity, NAFLD prevalence is high, which highlights a significant metabolic burden.(14,15) Both NAFLD and PCOS share a same pathophysiological basis. Insulin resistance is a key driver of NAFLD also, independent of obesity or diabetes, and plays a central role in hepatic fat accumulation by promoting increased lipogenesis and impaired fat oxidation(16). It has been reported that even in lean women with PCOS, insulin resistance is still a common feature in the Indian population(17). In these patients, the use of insulin sensitizers has been reported to improve menstrual cyclicity, induce ovulation, enhance fertility, and increase endometrial receptivity, thus creating a strong link between insulin resistance and female reproductive function(11,18). This presentation of NAFLD/ PCOS in non-obese patients highlights the importance of diet composition, genetic background, and timing of exposure in modulating disease phenotype.

Despite the increasing burden of metabolic and reproductive disorders in women, most of the DIO models are done in male mice, causing a gap in sex specific metabolic responses. C57BL/6J female mice are more resistant to diet induced weight gain, glucose intolerance and insulin resistant when compared to their male counterparts(19,20). PCOS in these models is commonly induced through pre- or post-natal exposure to DHEA or DHT, Letrozole, estradiol valerate and anti mullerian hormone(21,22). Though these models mimic PCOS features by imparting hyper androgenism and polycystic ovaries, the data on metabolic disturbances caused in these mice is inconclusive. A HFD or HFHS diet with Letrozole or DHEA dosing in female rodents disrupts the normal metabolic functioning in addition to induction of Polycystic ovaries with hyperandrogenism in them(23–25).

Current studies on PCOS or NA FLD rely upon prenatal or neonatal androgen exposure and early postnatal dietary exposure to induce disease phenotypes. This fails to address the adolescent onset metabolic disturbances often triggered by improper eating patterns and high fat and high sugar intake in humans.

In this study we investigated the differential effects of in house HFHS diet and HFD (60% fat) in 6 weeks old post pubertal C57BL/6J female mice which is the most critical period in reproductive maturation and significant metabolic changes. The findings of this study highlight the importance of diet composition and timing in modelling PCOS and NAFLD in female rodents.

## Materials and methods

### 1.1. Experimental mice and Diet

All procedures were performed with the approval of animal Ethics Committee of Indian Institute of Technology Madras (IITM/IAE/MDX-1/12/2021). C57BL/6J mice of four weeks age were procured and then acclimatized for 14 days in a temperature-controlled mice room with 12 hours light and dark cycle with ad-libitum access to water and food. At 6 weeks of age the mice from the first cohort were randomly assigned to Normal fat normal sugar (NFNS) group receiving standard chow diet and a High fat high sugar (HFHS) group receiving a homemade high fat and high sugar diet. The diet was prepared in house by combining beef lard and fructose to standard chow diet which mimics the western diet composing of 45% fat and 10% sugar (Figure 1 A). The second cohort of mice received a diet with 10 kcal % Fat (Cat# D12450Ji) for control group and 60 kcal % Fat (Cat# D12492i) for High fat diet group. The diets were purchased from research diets Inc. Both the cohorts had ad-libitum access to water and food and the diet was continued for twenty-one weeks following which mice were euthanized. The experimental time line is given in Figure 1 B. Body weight and abdominal circumference of the mice were monitored continuously.

**Figure 1.**
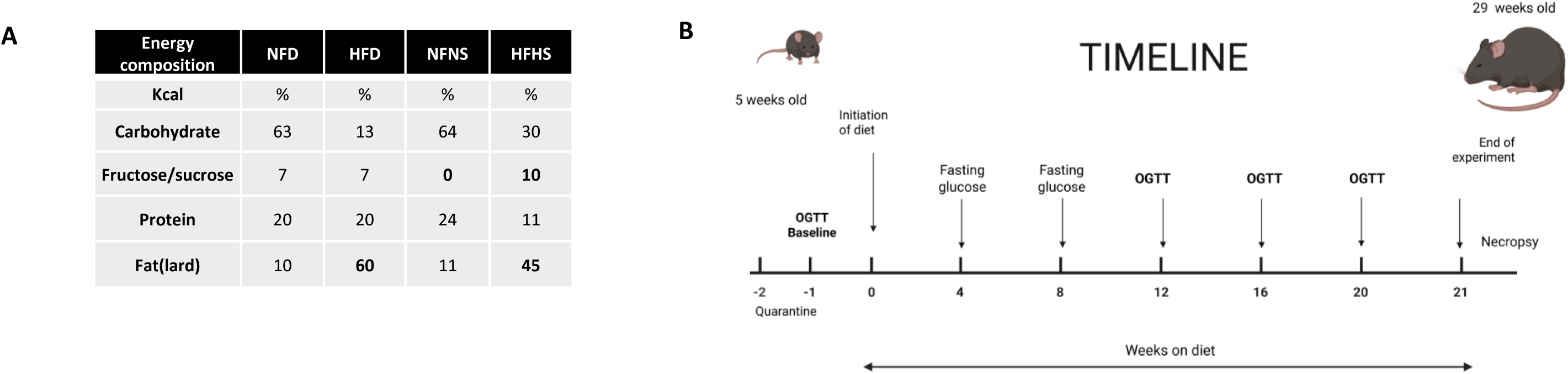
Experimental outline. (A) Macronutrient composition of the High-Fat High-Sugar (HFHS) and High-Fat Diet (HFD) regimens used in this study compared with their respective control diets. (B) Experimental timeline detailing the 21-week dietary intervention in female C57BL/6J mice, with assessments including body weight, OGTT, estrous cyclicity, and tissue collection at study termination.

### **1.2.** OGTT and Blood glucose measurements

Mice were fasted for 6 hours and 2gm/kg body weight glucose was administered orally by gavage. The measurements were done baseline and 12, 16, 20 weeks on diet. Blood was withdrawn through tail vein bleeding. Glucose (mg/dl) were measured at 0, 30, 60, and 120 minutes using Accu check glucometer. OGTT data was plotted as area under the curve (AUC).

### 1.3. Estrous cycle monitoring

To monitor the estrous cycle, vaginal smear was done every day for a period of 10 days starting 18 weeks of feeding in cohort 2.Vaginal swab was taken using cotton swab soaked in 1xPBS and then smeared on glass slides. Dried vaginal smear slides were dipped in 0.1% crystal violet solution for 3–5 min to stain the cells. The slides were subsequently washed with double-distilled water. After washing, a stained smear is fixed and mounted using a coverslip. The slides were immediately imaged using Leica upright microscope at 4X magnification. The estrous cycle stage was defined based on the cell types found in vaginal cytology. Nucleated epithelial cells represented the proestrus stage, and cornified squamous epithelial cells predominated in the estrus stage. The metestrus stage consisted of both cornified squamous epithelial cells and leukocytes, while the diestrus phase was noted for the predominance of leukocytes.

### 2.4 Tissue collection

After the diet treatment for 21 weeks, mice were sacrificed to harvest the tissues. Mice were fasted for 6 hours and euthanized using isoflurane inhalation. Blood was collected instantly via cardiac puncture, allowed time to clot, then centrifuged for 10 min at 1000g. Serum was stored at -80 °C until further usage. Mice were dissected and organs were excised carefully, washed with PBS and stored in 10% neutral buffered formalin for histopathology.

### 2.5 Histopathology of Ovarian and Liver tissues

Ten percent neutral-buffered formalin-fixed tissues were paraffin embedded, sectioned, and hematoxylin and eosin (H&E) stained. Hepatic fibrosis was assessed by Masson’s trichrome staining. Specimens were scored for the severity of hepatocellular steatosis, ballooning, inflammation and fibrosis by a pathologist. Ovarian antral follicles, corpora lutea and cystic follicles were assessed by H&E staining by a pathologist.

### 2.6 Biochemical measurements

Insulin measurement was done at the end of the study using Mercodia Mouse Insulin ELISA (Cat# 10-1247-01) as per the manufacturers protocol. Biochemical parameters such as lipid profile and liver enzymes were measured at the end of the study. Insulin resistance and β cell function were evaluated using the Homeostasis Model Assessment Method. The HOMA-IR and HOMA-β was calculated using the below mentioned following formula:

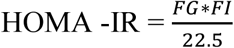

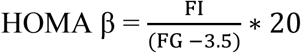

where FI is fasting insulin (in µU/mL) and FG is fasting glucose (in mmol/L).

### 2.6. Statistical analysis

Data are presented as mean ± SEM. Statistical analyses were performed with GraphPad Prism 8.0 software. Based on the experimental, one-way or two-way ANOVA was performed to analyse the data. A P value < 0.05 was considered for statistically significance.

## 2. Results

### 3.1 C57BL/6J female mice are resistant to gaining weight after fed with HFHS diet

Mice were fed with HFHS diet for continuously 21 weeks. To determine the effects of the HFHS diet on weight gain, animals were weighed on fasting for every four weeks until sacrifice. Mice were not showing any significant weight gain even upon feeding HFHS diet for 21 weeks (Figure 2A). There was no significant changes observed in the Fasting blood glucose (Figure 2 B). Interestingly these mice showed an increased abdominal circumference upon feeding HFHS for twenty one week which suggests abdominal adiposity and liver enlargement (Figure 2C).

**Figure 2:**
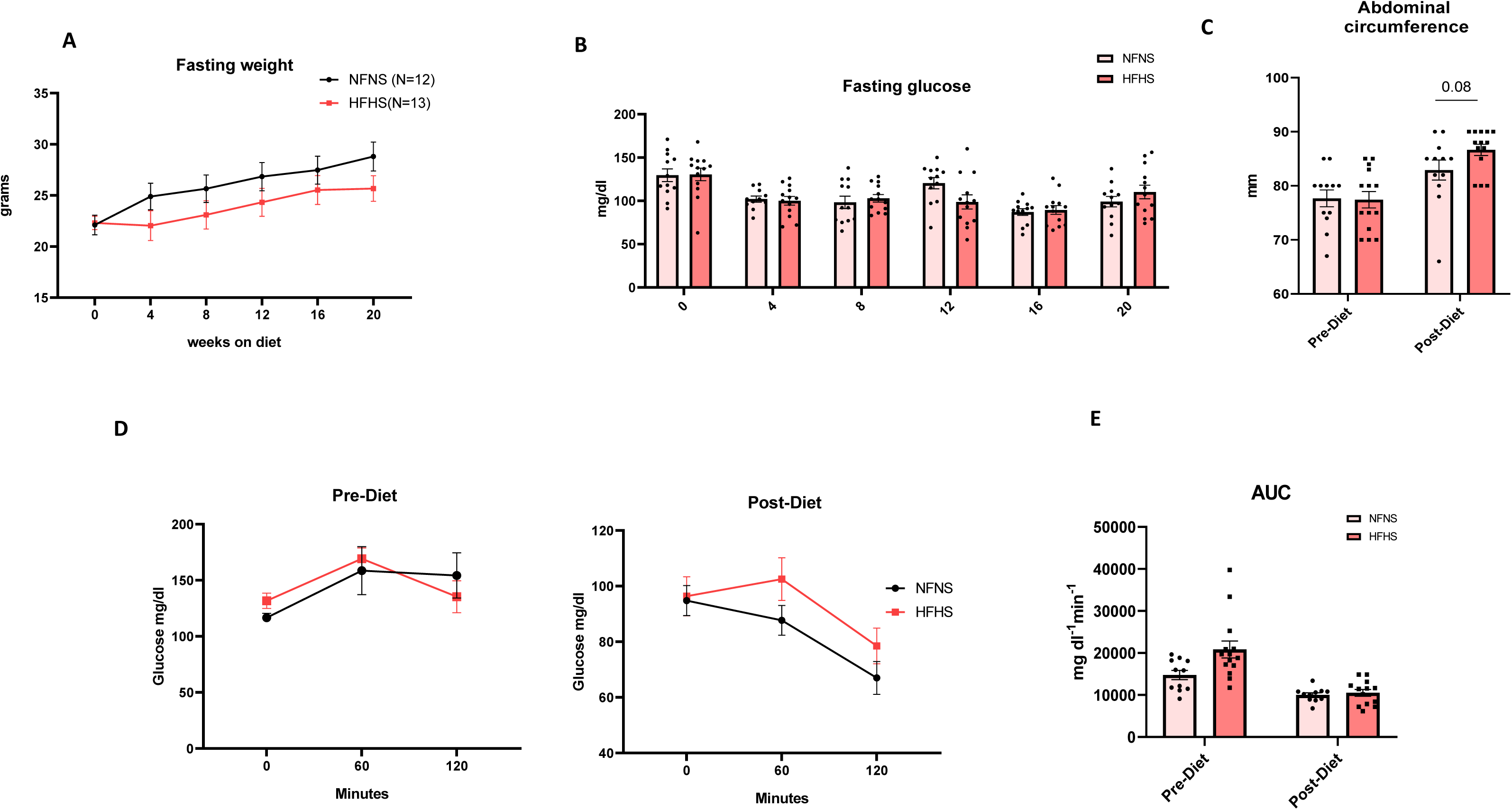
**HFHS feeding does not cause increase in body weight and alter glucose metabolism in C57BL/6J female mice** (A) Fasting weights and (B)Fasting glucose measured every week. (C) Abdominal circumference measured pre and post diet. (D) Blood glucose measurements (mg/dL) at 60-minute intervals after a 2 g/kg glucose injection of mice at 0 and 21 weeks on diet. (E) Respective area under the curve post and pre diet. (N=10-13 in each group). All the data points are represented as mean±sem. One way annova was used to compare the variances with a post hoc test. *\*P* < .05; *\*\*P* < .01; *\*\*\*P* < .001; *\*\*\*\*P* < .0001 was considered significance.

### 3.2. Female C57BL/6J mice are resistant to glucose intolerance despite feeding a HFHS diet

Oral Glucose Tolerance Test (OGTT) was performed in Cohort 1 every four weeks from baseline (week 0) to week 20 of HFHS feeding to assess the development of glucose intolerance. OGTT results revealed that these mice did not develop impaired glucose tolerance during the course of the study. A comparison of OGTT curves at baseline and at the end of 20 weeks showed no significant changes in the Fasting blood glucose and post load glucose levels (Figure 2D). The AUC for glucose clearance also did not change significantly throughout the period of diet feeding (Figure 2 E). Histology analysis of ovarian tissue showed none of the mice developed cystic follicles. (Supplementary Figure 1)

### 3.3 HFHS diet induces dyslipidaemia and Hepatic Steatosis in female C57BL/6J

Since we saw an increase in abdominal circumference and a fatty liver change during necropsy we sought to measure serum liver enzymes in these mice. The HFHS mice showed significantly higher levels of Serum Glutamate Oxaloacetate Transaminase (SGOT), Alkaline Phosphatase (ALP), and Gamma-Glut amyl Transferase (GGT) (Figure 3A). Serum lipid profile measurements showed that the HFHS mice developed dyslipidaemia. They had significantly higher levels of total cholesterol (TC), High density lipoprotein (HDL), Low density lipoprotein (LDL) when compared to control mice (Figure 3 b). In order to evaluate whether prolonged HFHS diet promotes NAFLD, we performed H&E staining of liver. Histopathological analysis of Liver tissue showed hepatocytes with Micro vesicular steatosis represented by multivacuolated cytoplasm and centrally placed nuclei, macrovesicular steatosis represented by macro vesicular large univacuolated cytoplasm with peripherally pushed nuclei in HFHS mice(Figure 4B, 4 D and 4 E). Masson’s trichrome staining showed portal fibrosis and fibrosis in the hepatic lobules (Figure 4C and 4 f). Mild to dense periportal inflammation was also observed (Figure 4 G). This suggests that HFHS diet promotes liver steatosis in C57BL/6J female mice.

**Figure 3:**
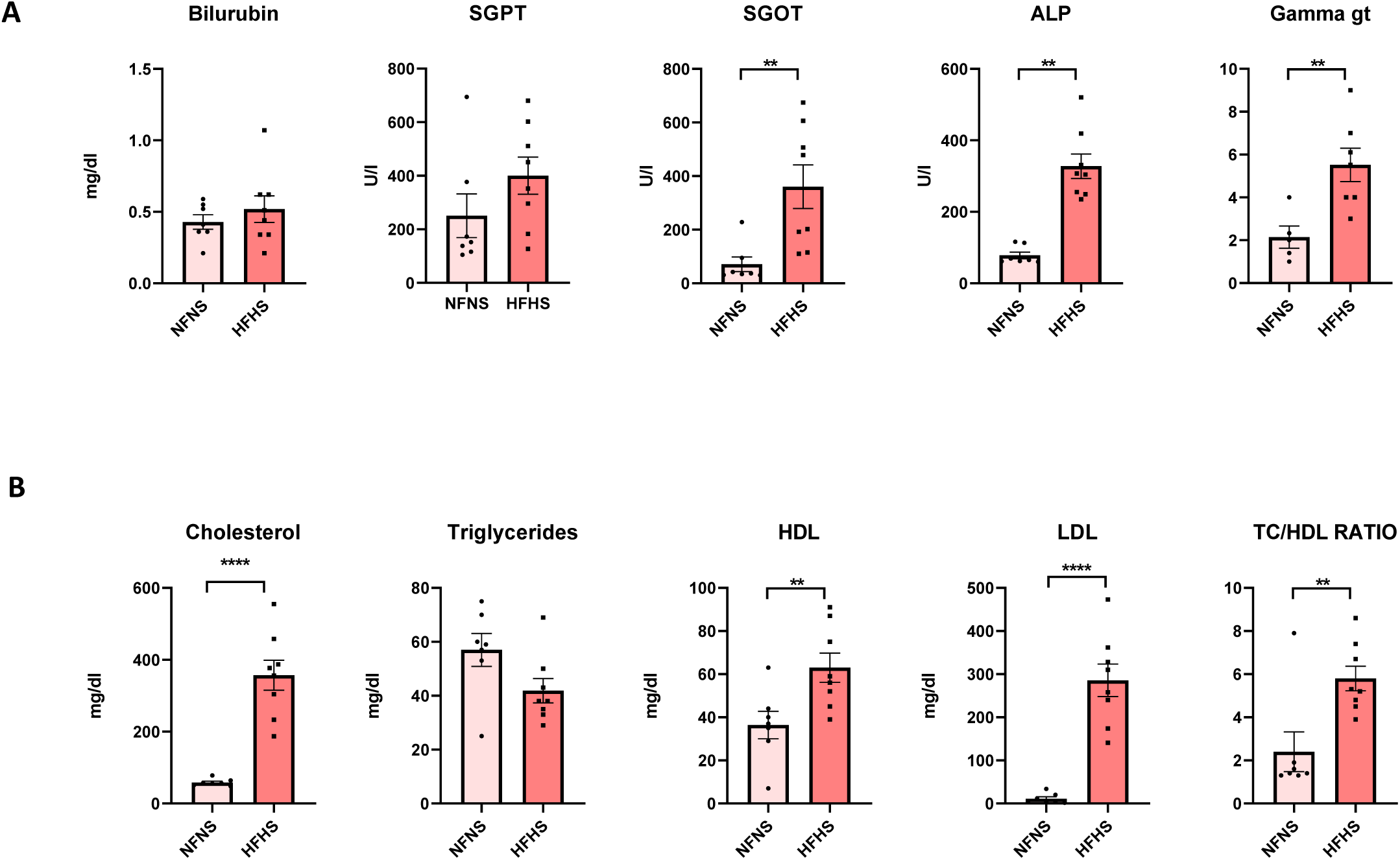
HFHS diet induces dyslipidemia and abnormal liver enzymes in female C57BL/6J. (A) Serum liver function test consisting if Bilirubin, SGPT, SGOT, ALP and gamma GT measure 21 weeks post HFHS diet. (B) Serum lipid profile measuring Cholesterol, Triglycerides, HDL, LDL and TC/HDL ratio. (n= minimum 5 in each group) Data are represented as Mean ±SEM. Unpaired T-Test was used to analyze the difference in mean of each category. *\*P* < .05; *\*\*P* < .01; *\*\*\*P* < .001; *\*\*\*\*P* < .0001.

**Figure 4:**
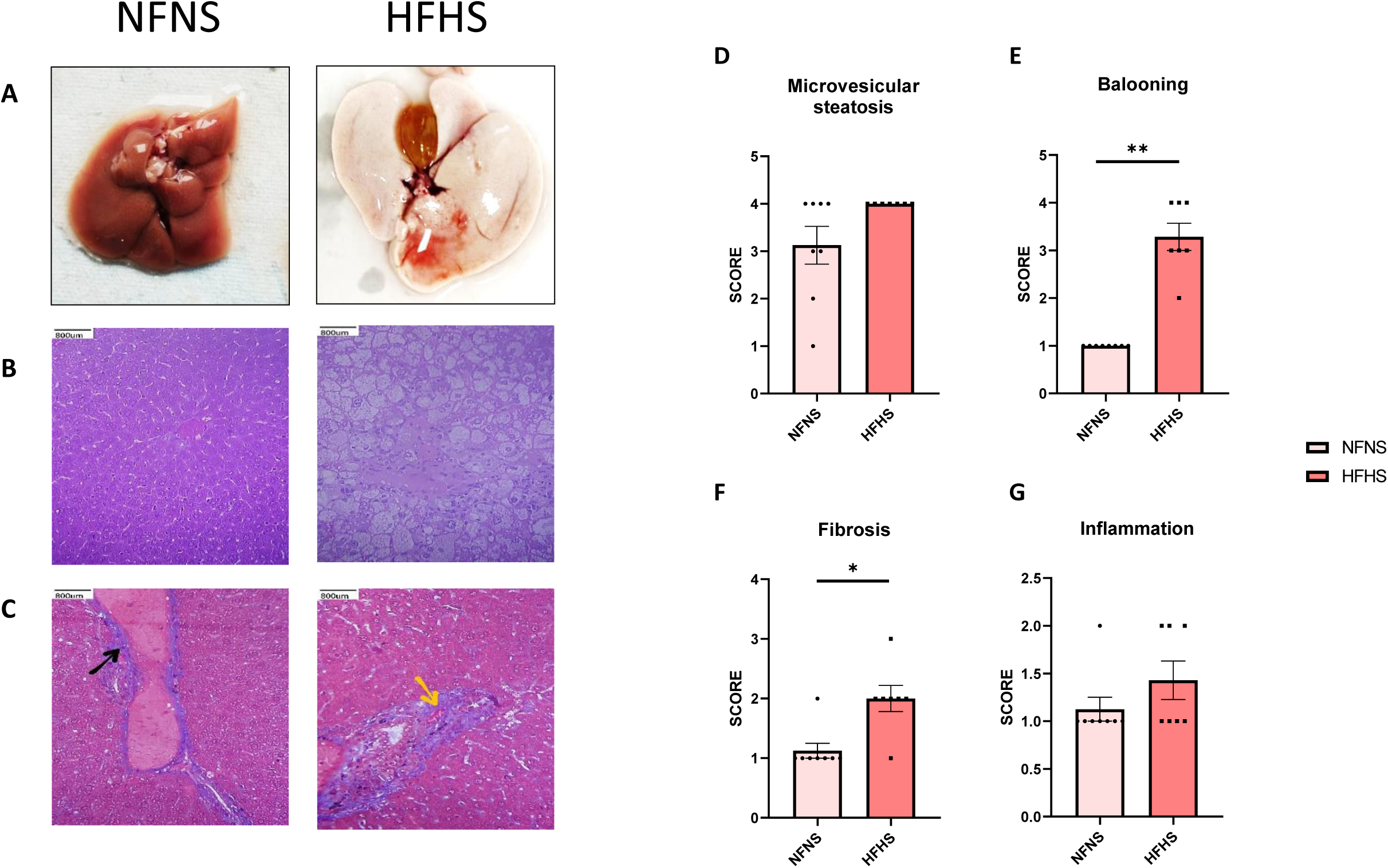
Histological assessment of the liver tissue in mice subjected to prolonged high fat high sugar-feeding shows severe hepatic steatosis. (A) Representative photographs of normal and fatty liver from NFNS and HFHS mice. (B) H&E staining of the liver sections. C) Masson’s trichrome staining of liver sections. Bar graph summarizing scoring of Microvesiclular steatosis(D), balooning (E), fibrosis (F) and inflammation (G) done by a pathologist. (N=6 animals minimum). Data are represented as mean±sem. Unpaired T-Test was used to analyze the difference in mean of each category. *\*P* < .05; *\*\*P* < .01; *\*\*\*P* < .001; *\*\*\*\*P* < .0001.

### 3.4 HFD induces metabolic dsyfunction in female C57BL/6J mice

Since we did not succeed in generating a PCOS phenotype with metabolic abnormalities using HFHS diet, we tried using a 60% HFD to induce PCOS like features in C57BL/6J mice. Upon feeding with HFD, these mice showed a marginal increase in weight but no significant difference in the abdominal circumference.

These mice showed impaired glucose tolerance from the twelfth week onwards. Due to a considerable variability in glucose clearance and weight in HFD mice, we tried to segregate them based on their diabetic phenotypes as mentioned in Burcelin et al (26). The AUCs were plotted against weight of each mice. HFD mice were considered obese or lean when their body weight was higher than the mean + 3 SD or lower than the mean + 1 SD of the NFD mice. Similarly, HFD mice were considered Diabetic or non-diabetic if their AUC was higher than mean + 3 SD or lower than mean + 1 SD. The ones in-between mean +1 SD and mean +3 SD were considered pre-Diabetic. There were four mice that are Diabetic and 4 mice that are prediabetic out of 16 HFD mice (Supplementary Figure 2). We considered all the eight as Diabetic phenotype and categorized them as HFD Diabetic (HFD DB). The mice below the mean +1 SD were categorized ad HFD non-Diabetic (HFD ND). None of the HFD mice were stratified as obese because these mice were not showing a significant change in body weight. One NFD mice that showed increased body weight and AUC was removed from the study.

HFD DB mice showed marginal increase in weight upon feeding with HFD for 21 weeks (figure 5 a). These mice also showed impaired glucose tolerance starting from twelfth week till twenty weeks (figure 5 B). These mice expressed a significantly decreased glucose clearance as seen in AUC graph (Figure 5 B and 5C). These HFD DB mice showed hyperglycaemia at the end of the die (Figure 6A). These mice also showed higher fasting insulin in terminal measurements. They were also insulin resistant and showed lower beta cell function (Figure 6A).

**Figure 5:**
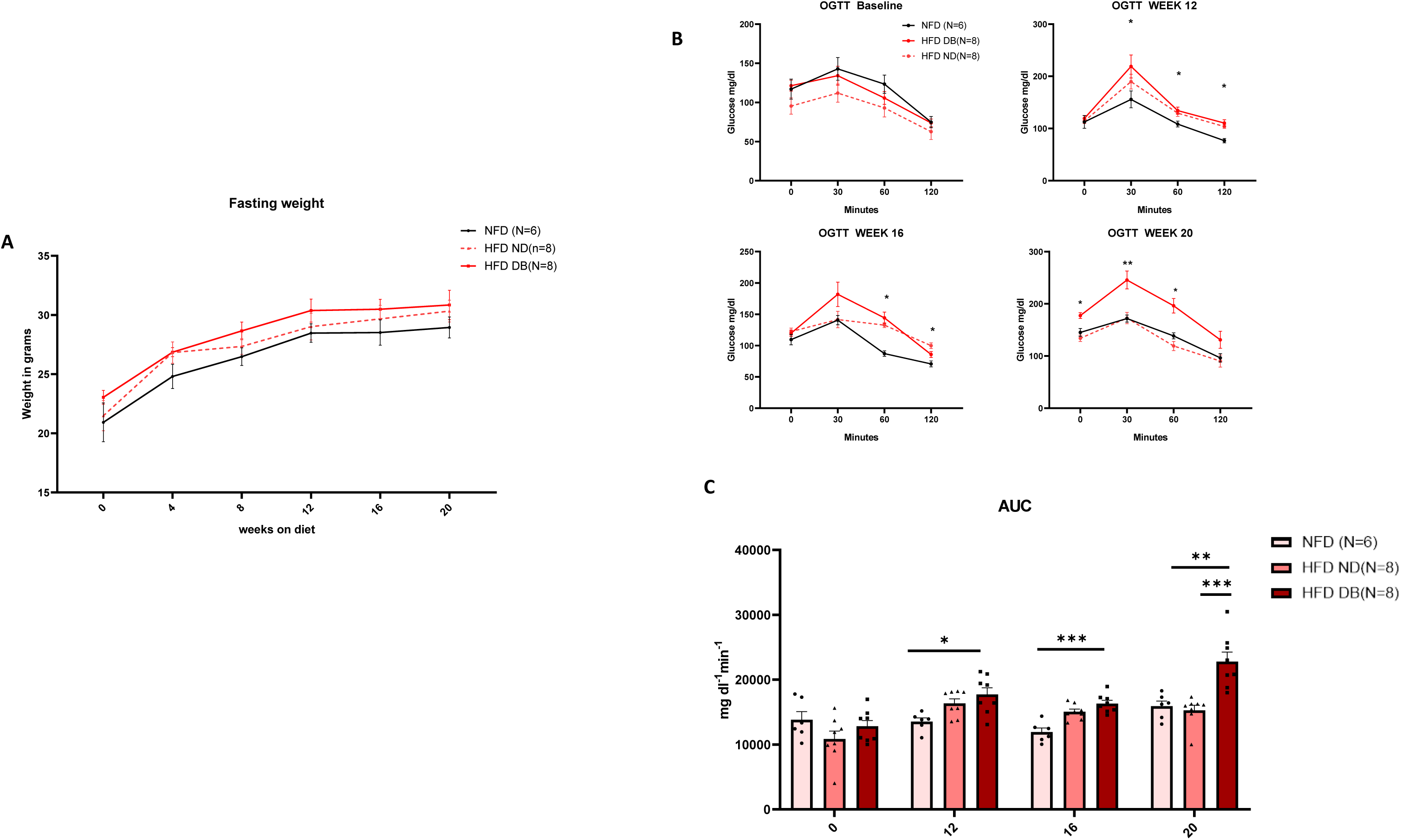
Effect of 60% HFD on weight gain and glucose clearance in C57BL/6J mice. (a) Fasting weight measured every four weeks. (b) oral glucose tolerance test done every four weeks until 20^th^ week. (c) Area under the curve showing glucose clearance. Data are represented as mean±sem. Unpaired T-Test was used to analyze the difference in mean of each category. *\*P* < .05; *\*\*P* < .01; *\*\*\*P* < .001; *\*\*\*\*P* < .0001.

**Figure 6:**
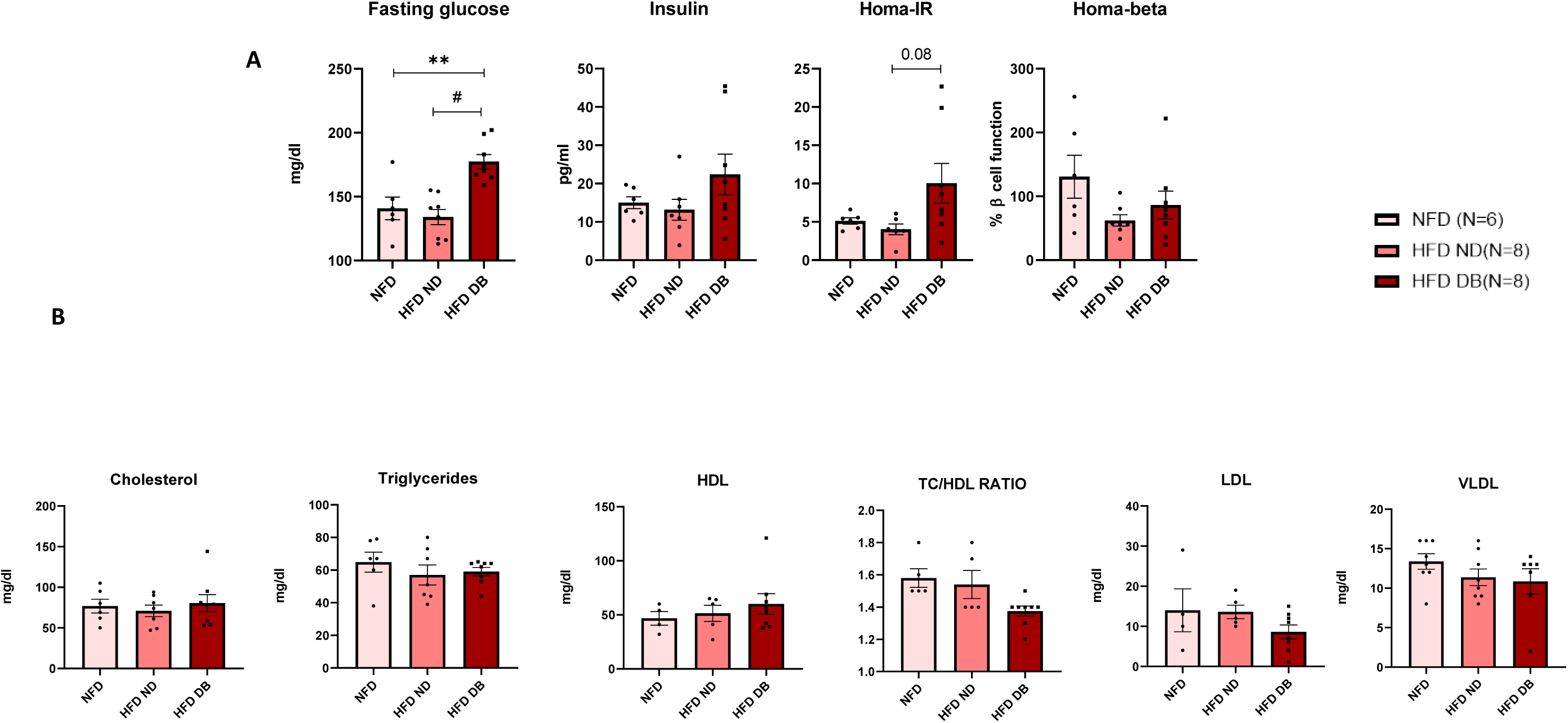
Effect of High fat diet on serum insulin and lipid profile in C57BL/6J female mice. (A) bar graph summarizing serum insulin levels and glucose levels and empirical HOMA indices calculation after 21 weeks of high fat diet. (B) Bar graph summarizing serum lipid profile after 21 weeks of high fat. Data are represented as mean±sem. Unpaired T-Test was used to analyze the difference in mean of each category. *\*P* < .05; *\*\*P* < .01; *\*\*\*P* < .001; *\*\*\*\*P* < .0001.

### 3.5 Female C57BL/6J mice are resistant to NAFLD development on HFD alone

Even upon feeding HFD for 21 weeks, none of the mice developed dyslipidaemia (Figure 6 B). The abdominal circumference was also not significantly different. In contrast to the steatotic changes observed in HFHS-fed mice, livers from mice fed the 60% high-fat diet showed minimal histopathological evidence of fatty liver disease. H&E staining revealed largely intact hepatic architecture with no significant micro vesicular or macrovesicular steatosis and ballooning. Periportal or lobular inflammation was either absent or mild. Masson’s trichrome stain also showed no significant fibrosis. (Supplementary Figure 3)

### 3.6. HFD induces PCOS like ovarian morphology in female C57BL/6J

Menstrual disturbance is a typical clinical feature of PCOS women. In this study all of the control mice had a normal estrous cyclicity indicating a healthy reproductive function. In contrast, mice fed with a high fat diet irrespective of the diabetic characterization showed clear disruption of estrous cyclicity (Figure 7C). HFD mice predominantly remained in the estrous phase for extended durations, indicating a disruption of normal cyclicity and suggesting that their reproductive cycle is arrested in the follicular phase (Figure 7 D). Cystic follicles a hallmark feature of PCOS were not present in control mice, whereas the HFD DB group showed significantly increased number of cystic follicles reflecting impaired follicular development characteristic of PCOS. In addition to it HFD mice also exhibited significantly lower number of corpora lutea (Figure 7 B).

**Figure 7:**
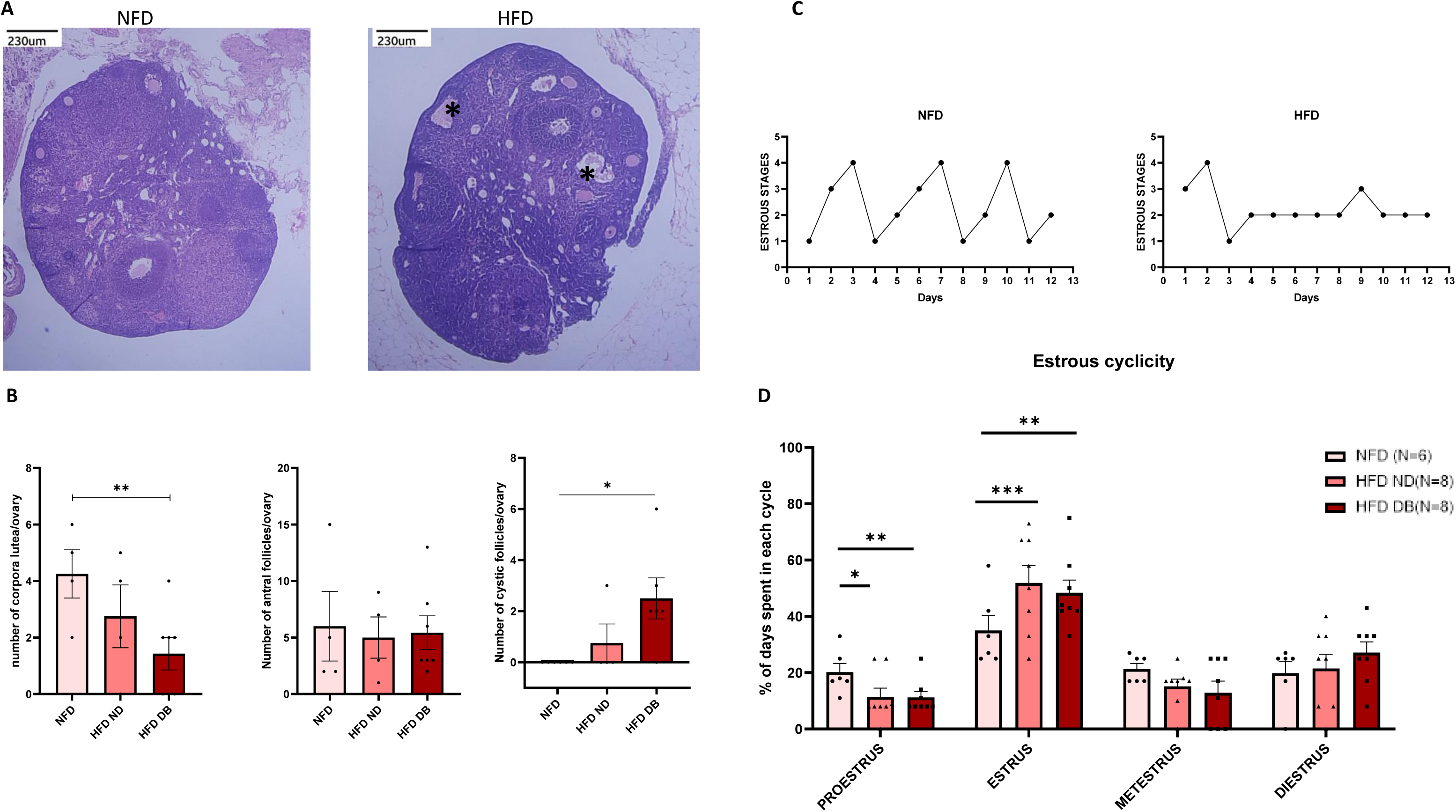
**Effect of HFD on estrous cyclicity and ovarian morphology in C57BL/6J female mice** (A) H&E staining of mouse ovarian tissue from NFD mice and HFD mice. Asterisk indicates cystic follicles. (B) Bar graph summarizing number of corpora lutea, antral follices and cystic follicles per ovary/mice. (C) Estrous cycle graph of NFD mice and HFD mice. (D) bar graph summarizing percentage of time, the mice had spent in each estrous cycle phase across a period of 12 days.

### 3.7. Correlation analysis

A correlation analysis was performed on biochemical parameters and cysts development and HFD phenotype. Spearman correlation analysis of Disease state (coded as NFD < HFD ND < HFD DB) showed significant positive correlations for Cystic follicles (ρ = 0.5305), Fasting glucose (ρ = 0.3766), Fasting insulin (ρ = 0.3633), HOMA-IR (ρ = 0.3814) Glucose intolerance (ρ = 0.6491). Presence of Cystic follicles was also significantly correlated with Fasting glucose (ρ = 0.6396), HOMA-IR (ρ = 0.6305) and glucose intolerance (ρ = 0.6867) suggesting a glucose centric influence of ovarian function. These findings indicate that progression from NFD to HFD ND and HFD DB leads to worsening of glucose homeostasis, hyperinsulinemia, and reproductive pathology.

## 3. Discussion

This study aimed to explore the differential effects of two obesogenic diets namely High-Fat High-Sugar (HFHS) and High-Fat Diet (HFD, 60%) on the development NAFLD and PCOS in post–pubertal female C57BL/6J mice. Our findings reveal that the composition of diet plays a crucial role in determining tissue-specific manifestation of metabolic disorders, with the HFHS diet predominantly inducing hepatic changes characteristic of NAFLD, while the 60% HFD preferentially triggered reproductive abnormalities consistent with PCOS.

One of the key findings from our study is the resistance of female C57BL/6J mice to glucose intolerance and diet induced obesity despite feeding for twenty one weeks on HFHS diet. This confirms the earlier studies done on sex specific responses of HFHS diet in C57BL/6J mice, where female mice are shown to be more resistant to develop metabolic syndrome than their male counterparts(20,27,28). This resistance may be attributed by the estrogen mediated protection against metabolic stress. Although these mice developed abdominal adiposity and dyslipidemia, their fasting glucose and AUC upon OGTT remained same. This suggests that abdominal fat accumulation need not induce abnormalities in glucose metabolism in this strain upon HFHS feeding. Histopathology analysis of liver showed hepatic steatosis including micro and macro vascular steatosis indicating that hepatic dysfunction and steatosis can progress independently of obesity or glucose intolerance, especially in female mice. These mice also did not develop Polycystic ovarian changes as seen in the H&E staining of tissue section of ovarian tissues despite feeding HFHS diet which is on contrary known to induce PCOS features upon feeding with postnatal hormone treated mice(24,25). In the absence of such hormonal priming, diet alone which also failed to induce metabolic alterations of glucose may not be sufficient to induce reproductive endocrine dysfunction or structural ovarian changes, suggesting that glucose metabolism may play a critical permissive role in the manifestation of PCOS features.

A 60% high fat diet in contrast induced systemic insulin resistance and glucose intolerance in these mice confirming previous results obtained from other studys(29). These mice upon feeding on HFD diet for 21 weeks showed a significant metabolic response. The stratification of these mice based on Burcelin etal, emphasize the heterogeneity in response to dietary fat exposure, which is a very common problem when involving rodent model systems(26).

Unlike the HFHS group, mice fed a high-fat diet that developed metabolic dysfunction (HFD DB group) demonstrated significant impairments in glucose clearance and insulin sensitivity, alongside the deveopment of PCOS-like ovarian pathology. The presence of cystic follicles and disrupted folliculogenesis in these HFD DB animals highlights the critical role of glucose metabolism in driving ovarian dysfunction. This was further validated by Spearman correlation analysis, where disease severity, progressing from NFD to HFD ND and HFD DB positively correlated with key metabolic and reproductive parameters. The presence of cystic follicle showed a strong positive correlations with fasting glucose (ρ = 0.6396), HOMA-IR (ρ = 0.6305), and glucose intolerance (ρ = 0.6867), pointing towards a glucose-centric model of reproductive impairment. These findings backs up the hypothesis that metabolic disturbances, particularly hyperglycemia and insulin resistance, are closely intertwined with ovarian pathophysiology in PCOS(30,31). We also found a strong correlation between disease progression and parameters such as glucose intolerance (ρ = 0.6491) and cystic follicle count (ρ = 0.5305) which suggests that a deteriorating metabolic state directly influences reproductive outcomes. Previous studies have similarly shown that HFD models, particularly when combined with perinatal or adult-onset insulin resistance, can induce PCOS-like features, including anovulation and cystic ovaries(23,24). This observed association between insulin resistance and ovarian morphology emphasizes the perception that hyperinsulinemia may intensify ovarian dysfunction and disrupt normal follicular maturation(9,32).

Together, these discoveries highlight that ovarian dysfunction in the HFD model is not only a consequence of dietary fat intake but is significantly potentiated by systemic metabolic derangements, supporting the need to consider metabolic health as a vital axis in the pathogenesis of PCOS.

The major limitation of our study was due to limited blood volume, we could not assess circulating sex hormone levels (e.g., testosterone, estradiol, LH/FSH ratios), which are crucial in defining PCOS phenotypes. Although the estrous cycling in rodents different from the human menstrual cycle in periodicity and hormonal profiles, limits its translational potential, the consistent development of PCOS-like ovarian morphology and glucose intolerance in the HFD-fed mice supports the utility of this model in studying diet-induced reproductive dysfunction. Future studies should include longitudinal hormonal profiling, adipose tissue cytokine analysis, and ovarian angiogenesis markers, which are known to be disrupted in both PCOS and NAFLD

Our study provides a translationally relevant model to explore PCOS and NAFLD as distinct but overlapping conditions. Our study draws several key conclusions. Firstly not all obesogenic diets induce PCOS or NAFLD in mice by disrupting the metabolism. HFHS diet induced hepatic and lipid disturbances without changing the metabolism and ovarian function but the HFD deregulated the glucose metabolism and reproductive function without having much effect on the hepatic axis. The heterogeneity in dietary exposures among human populations may account for variable expression of liver versus ovarian pathologies. Our findings highlight the need for tailored dietary interventions based on individual metabolic phenotypes.

---

**Supplementary figure 1:**
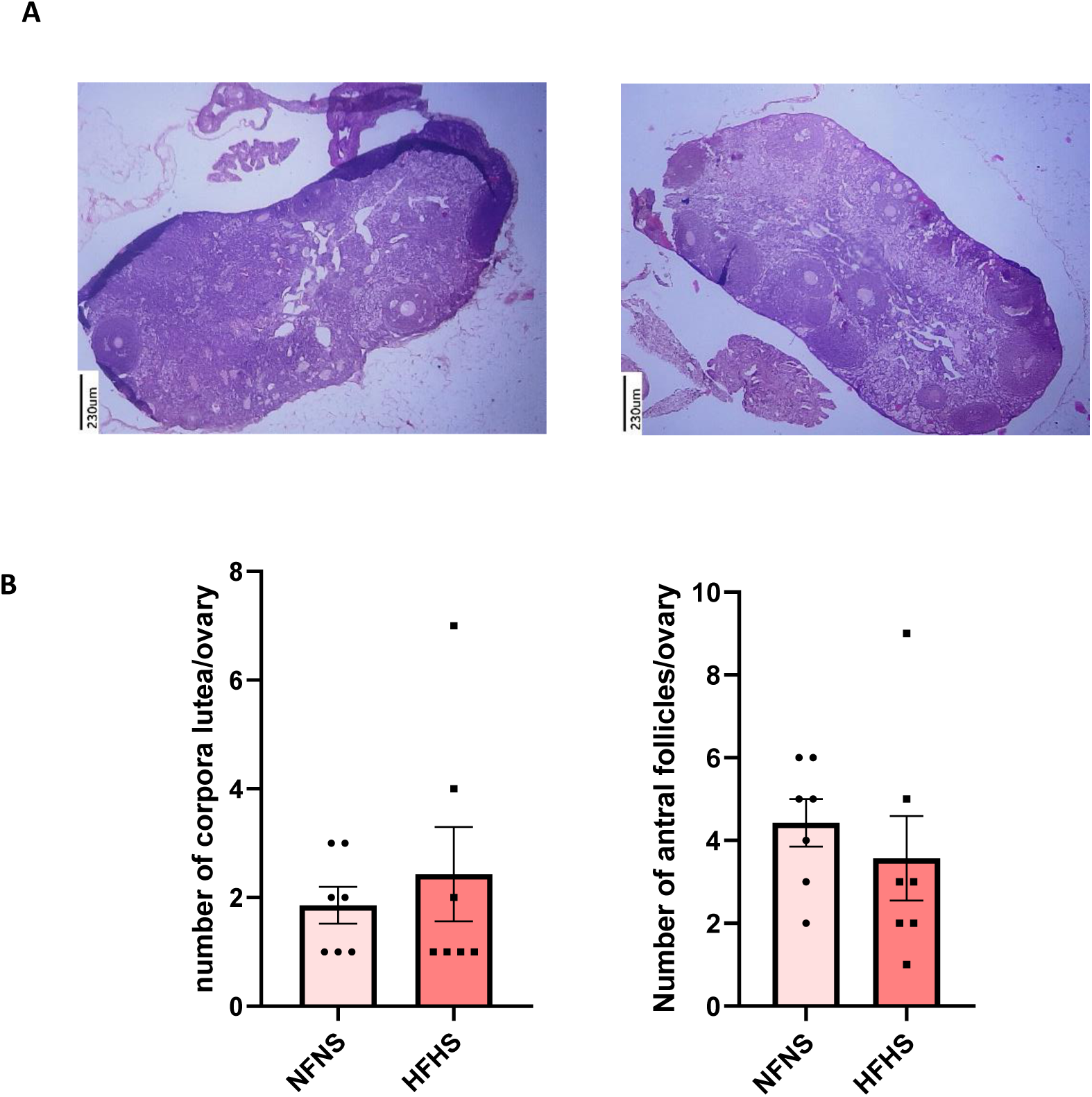
Histological assessment of ovary in mice subjected to prolonged high fat high sugar-feeding A) H&E staining of mouse ovarian tissue from NFD mice and HFD mice. (B) Bar graph summarizing number of corpora lutea, antral follices per ovary/mice.

**Supplementary figure 2 :**
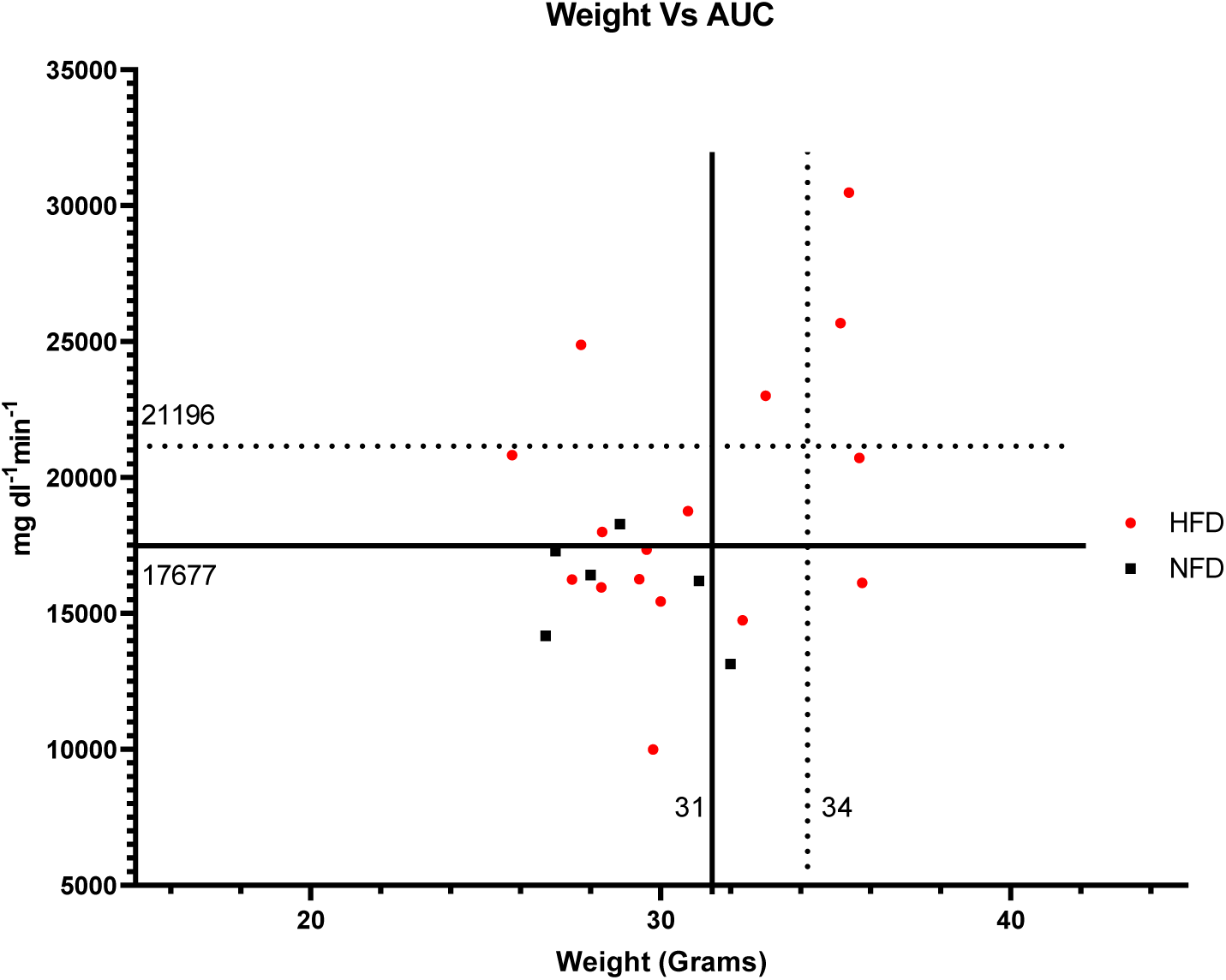
**Categorization of HFD mice based on metabolic parameters**. (A)Correlation of weight Vs AUC of the OGTT done at the end of the HFD study.

**Supplementary figure 3:**
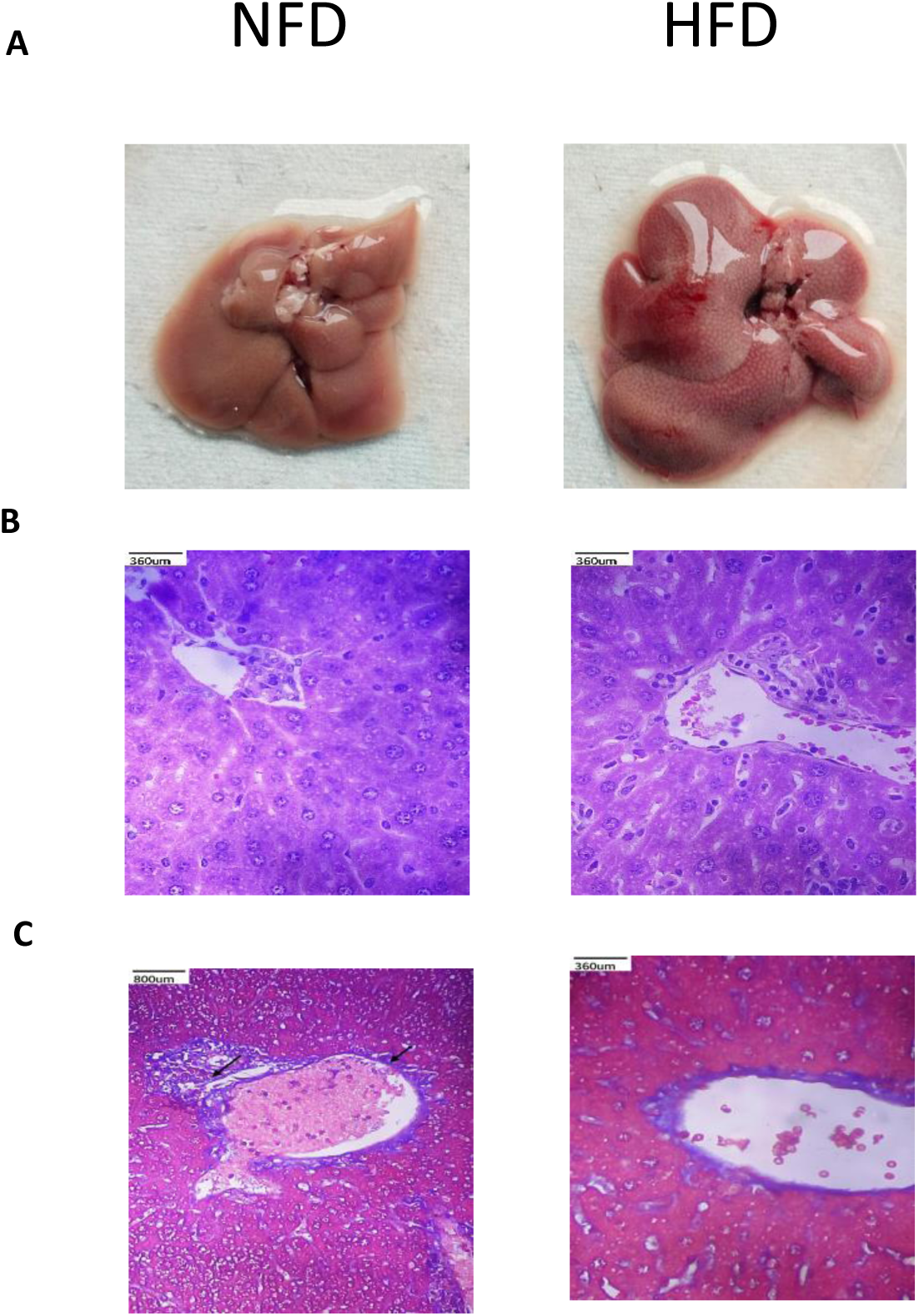
Histological assessment of liver in mice subjected to prolonged high fat feeding (A) Liver tissue from NFD and HFD mice. H&E(B) and Masson’s trichrome staining (C) of liver of the HFD and NFD mice.

